# A brain-enriched circRNA blood biomarker can predict response to SSRI antidepressants

**DOI:** 10.1101/2024.04.30.591973

**Authors:** Grigorios Papageorgiou, El-Cherif Ibrahim, Gabriella Maxson, Victor Gorgievski, Evelyn Lozano, Raoul Belzeaux, Thomas Carmody, Eleni T. Tzavara, Madhukar H. Trivedi, Nikolaos Mellios

**Affiliations:** Circular Genomics Inc, Albuquerque, NM; Previously at: University of New Mexico, Department of Neurosciences, Albuquerque, NM; Aix-Marseille Univ, CNRS, INT, Inst Neurosci Timone, Marseille, France; Université Paris Cité, CNRS, Integrative Neuroscience and Cognition Center, 75006 Paris, France; Hôpital Sainte Marguerite AP-HM, Pôle de Psychiatrie, 13274 Marseille, France; University of Texas Southwestern Medical Center, Department of Psychiatry, Dallas, Texas

## Abstract

Major Depressive Disorder (MDD) is a debilitating psychiatric disorder that currently affects more than 20% of the adult US population and is a leading cause of disability worldwide. Although treatment with antidepressants, such as Selective Serotonin Reuptake Inhibitors (SSRIs), has demonstrated clinical efficacy, the inherent complexity and heterogeneity of the disease and the “trial and error” approach in choosing the most effective antidepressant treatment for each patient, allows for only a subset of patients to achieve response to the first line of treatment. Circular RNAs (circRNAs), are highly stable and brain-enriched non-coding RNAs that are mainly derived from the backsplicing and covalent joining of exons and introns of protein-coding genes. They are known to be important for brain development and function, to cross the blood-brain-barrier, and to be highly sensitive to changes in neuronal activity or activation of various neuronal receptors. Here we present evidence of a brain-enriched circRNA that is regulated by Serotonin 5-HT2A and Brain-Derived Neurotrophic Factor (BDNF) receptor activity and whose expression in the blood can predict response to SSRI treatment. We present data using circRNA-specific PCR in baseline whole blood samples from the Establishing moderators and biosignatures of antidepressant response in clinical care (EMBARC) study, showing that before treatment this circRNA is differentially expressed between future responders and non-responders to sertraline. We further show that the expression of this circRNA is upregulated following sertraline treatment and that its trajectory of change post-treatment is associated with long-term remission. Furthermore, we show that the biomarker potential of this circRNA is specific to SSRIs, and not associated with prediction of response or remission after Placebo or Bupropion treatment. Lastly, we provide evidence in animal mechanistic and neuronal culture studies, suggesting that the same circRNA is enriched in the brain and is regulated by 5-HT2A and BDNF receptor signaling. Taken together, our data identify a brain-enriched circRNA associated with known mechanisms of antidepressant response that can serve as a blood biomarker for predicting response and remission with SSRI treatment.

## Introduction

The prevalence of Major Depressive Disorder (MDD) has been steadily increasing over the last few decades,^1^ but has, since the pandemic, reached epidemic proportions within the US, with close to 30% of the adult US population now reporting to have been diagnosed with depression during their lifetime.^2,3^ Even though multiple classes of antidepressants, such as selective serotonin reuptake inhibitors (SSRIs), serotonin and norepinephrine reuptake inhibitors (SNRIs), and norepinephrine dopamine reuptake inhibitors (NDRIs), have demonstrated significant efficacy and safety, they appear to only work for a subset of patients.^4,5^ As a result, less than 50% of patients with MDD respond to their first antidepressant treatment with remission rates being less than 30%.^4,5^ This trial-and-error approach leads to significant delays in properly managing MDD, imposing a notable burden to patients suffering with this devastating psychiatric disorder and resulting in significant costs to our healthcare system. Thus, there is a significant unmet need for developing reliable and robust biological biomarkers that could predict response to different antidepressants, thereby allowing for a precision medical approach to depression treatment.

Currently, the only widely available precision medicine tools to patients with MDD are associated with DNA-based pharmacogenomics, which are designed to predict an individual’s genetic predisposition for metabolizing different psychiatric drugs, including antidepressants.^6^ Although such approaches are aimed at the right direction of providing potential personalized information about treatment options, they are not able to achieve a direct prediction of response to antidepressant treatment. A known limitation of utilizing such genetic-based predictive biomarker approaches is the static nature of our genome versus the dynamic and adaptable nature of transcriptomic and proteomic biological signatures. However, the potential of any peripheral RNA and protein biomarker signatures to be capable of reliably predicting response to psychiatric drug treatments is significantly constrained by the fact that the vast majority of such molecules do not cross the blood-brain barrier (BBB) and are, thus, not directly associated with the molecular mechanisms that underlie psychiatric drug treatment response within the brain.

Circular RNAs (circRNAs) are a subtype of non-coding RNAs that are generated from non-canonical backsplicing and covalent joining of RNA, including exons and introns of protein-coding genes.^7–10^ They are the only type of RNA that has been found to be particularly enriched in the brain vs all other tissues and to be able to cross the BBB and to be readily-detectable in the blood.^7–9,11–15^ Due to their unique structure, they are particularly resistant to exonuclease degradation, and are thus known to have robust inherent stability and much longer half lives compared to linear mRNA molecules.^14,15^ CircRNAs are known to exert significant and diverse functional effects on gene expression by either sequestering microRNAs (miRNAs) and RNA-binding proteins (RBPs) or directly interacting with numerous RNA and protein partners to exert both transcriptional and post-transcriptional influence on a significant number of downstream gene targets.^16^ Not surprisingly, given their abundance in neural tissue and significant upstream impact on gene expression, circRNAs, have been shown to be important for brain development, maturation, and function, and have been linked to a plethora of psychiatric and neurological disorders.^11,13,17–28^ Due to the above characteristics, circRNAs, thus, appear to hold significant potential to become ideal molecular biomarkers for better diagnosis and treatment of psychiatric disorders.^29,30^

Here we provide evidence of a brain-enriched circRNA, whose expression in whole blood is significantly associated with response to sertraline treatment in a large well-characterized clinical biomarker study of antidepressant response. We highlight the specific nature of this circRNA biomarker at predicting response to sertraline treatment in baseline whole blood samples of patients with MDD, and uncover a dynamic expression profile post treatment that is associated with possibility of remission. In parallel, we show data of potential utility of the same circRNA biomarker in specifically predicting remission with SSRI class of antidepressants in cellular extracts from whole blood of patients with MDD in a smaller naturalistic antidepressant response study. Furthermore, we show that the expression of this circRNA in mouse brain is regulated by serotonin and Brain-derived neurotrophic factor (BDNF) receptor activity with the extracellular signal-regulated kinase (ERK) and cAMP response element-binding protein (CREB) signaling pathways being part of the downstream molecular cascades associated with the expression of this circRNA within neurons. Taken together, our data provide evidence of a biomarker that is predictive of response and remission after SSRI antidepressant treatment and is significantly associated with known molecular mechanisms of antidepressant response.

## Materials and Methods

### RNA extraction from human blood samples from EMBARC clinical study

We utilized samples and clinical information from the multi-site NIMH-funded Establishing Moderators and Biosignatures of Antidepressant Response in Clinical Care (EMBARC) study. This study was specifically designed to uncover biomarkers for prediction of response to antidepressant treatment. We separated 91 baseline PaxGene whole blood samples from patients treated with Sertraline randomly to a discovery (N = 49) and replication (N = 42) group. RNA isolation from human whole blood samples was done using the PaxGene Blood RNA kit 50, v2 IVD, PreAnalytiX (Qiagen, Hilden, Germany) following the manufacturer’s supplied protocol with few slight modifications. RNA quality (A260/280 and A260/230) as well as concentration of isolated total RNA was assayed through NanoDrop One Spectrophotometer (Thermo Fisher Scientific, Waltham, MA).

### RNA cleanup

Following RNA extraction, RNA was further processed using the Monarch RNA cleanup Kit (New England Biolabs, Ipswich, MA). Protocol steps were executed according to manufacturer’s instructions with slight modifications and the RNA was finally eluted in a total volume of 40ul of RNase/DNase free water. RNA quality and concentration were then again assayed using NanoDrop One Spectrophotometer (Thermo Fisher Scientific).

### Reverse transcription and circRNA-specific PCR.

Reverse transcription of 500ng of total RNA was performed using the SuperScript IV First-Strand Synthesis System (Thermo Fisher Scientific) per the manufacturer’s instruction and via use of random hexamers. Reverse transcription was performed using the Veriti Dx 384 well Thermal Cycler, designed for clinical use (Applied Biosystems, by Thermo Fischer Scientific). Quantitative real time PCR (qRT-PCR) was done using PowerUp SYBR Green Master Mix (Thermo Fisher Scientific) along with custom designed, validated, and sequence-verified circRNA primers. All circRNA qRT-PCR products were run on an agarose gel and sequence validated. At the end of each qPCR amplifications plots and melt curves (ΔRn vs cycle per well) were automatically calculated by the Quant Studio 7 Pro qPCR instrument (Thermo Fisher Scientific). Relative circRNA levels were quantified directly from the Ct average’s as shown before.^17,20^

### Venous blood leucocyte extraction and RNA isolation

Eligible study participants were part of the replication cohort of a larger multi-site naturalistic study of adults diagnosed with MDD (ClinicalTrials.gov with ID: NCT02209142). ^32^ 8-9ml of whole blood was collected in EDTA tubes and processed as described before. ^32^ Leukocytes were captured using the LeukoLOCKTM filter (Thermo Fisher Scientific). Briefly, the filter with the captured leucokytes was washed with RNAlater® and then moved to a –80°C freezer. TRI reagent (Thermo Fisher Scientific) was used to lyse the cells, which were then mixed with Bromo-3-chloro-propane (Sigma Aldrich, St. Louis, MO). Following centrifugation, total RNA was precipitated with ethanol and further purified and washed 0.1mM EDTA. Total RNA was then treated with DNase using the DNA-freeTM kit (Thermo Fisher Scientific).

### RNA extraction and circRNA quantification in cell cultures and mouse brain tissue

For RNA isolation of cultured cells or brain tissues extracted from mouse brains, RNA was isolated using the miRNeasy RNA isolation kit (Qiagen) following the manufacturer’s supplied protocol. RNA quality as well as concentration of isolated total RNA was assayed through Nanodrop 2000 spectrophotometer (Thermo Fisher Scientific), with all samples passing the quality control measurements (A260/230 and A260/280). Reverse transcription of total RNA (100ng or 500ng) and circRNA-specific PCR was done as described above.^17,20^

### SH-SY5y Human neuroblastoma cell line treatments

The SH-SY5Y epithelial human neuroblastoma cell line was purchased from ATCC (Manassas, VA; product code CRL-2266™). Their morphology and viability were observed daily under the microscope. After reaching passage #7, SY-5 cells were plated in a 24-well plate at a concentration of 100,000 cells per well. 48 hours after been plated, SH-SY5 cells were treated with ANA-12, an industrial TrkB receptor antagonist, purchased from Tocris Bioscience (Bristol, United Kingdom; catalog no 4781). ANA-12 was dissolved in DMSO and to compensate for the effects of DMSO in cells viability, a negative control / vehicle group was included with the same concentration of DMSO. 24 hours after treatment, cells were harvested, and processed as described above using the miRNeasy RNA isolation kit (Qiagen).

### Primary mouse cortical neuron pharmacological treatments

Mouse cortical neuronal cultures were purchased from Thermo Fisher Scientific – Catalog number: A15585. Neurons were isolated and cryopreserved from C57BL/6 mice at embryonic day-17. Neurons were plated at a density of 4×10^4 cells/12-mm at a coverslip coated with poly-Ornithine on a 24-well plate. Neurons were allowed to adhere for about 30 min before adding 500uL of their respective media. Neurons were fed by replacing half the plating media volume with fresh media every third day. A media of a combination of Neurobasal Plus, 1XB27 Plus, 2mM Glutamax and 5% Pen/Strep was used (all items purchased from Thermo Fisher Scientific). At DIV13, when neurons reach their maturation state (as observed with microscopy), pharmacological treatments were conducted as mentioned below. All the small molecule inhibitors were dissolved in DMSO. A negative control / vehicle group was included with the same concentration of DMSO. 24 hours after treatment, neurons were subjected to RNA extraction as shown above.

CREB and CREB-CBP pharmacological inhibition: For restricting the interaction of CREB with its co-transcriptional activator CBP, a specific KIX-KID interaction inhibitor was used (CAS 92-78-4; Millipore/Sigma Aldrich) was added at a concertation of 2.5µM.

Small molecule kinase inhibitors: To examine which intracellular neuronal induced kinase can alter the levels of our circRNA of interest, various candidates were tested in primary cortical neurons. All pharmacological agents were added at DIV13 (dissolved in DMSO) and were purchased from Tocris Bioscience corporation. H 89 dihydrochloride (Cat. No. 2910), a selective PKA inhibitor was added at a dose equal to 120nM. GF 109203X (Cat. No. 0741) was used to inhibit PKC activity at a dose equal to 0.2µM. FR 180204 (Cat. No. 3706) was used for ERK1,2 inhibition at a dose of 0.3 µM.

### In vivo Pharmacological treatments

#### Animals

All animal protocols complied with French and European Ethical regulations. The experimental protocols were approved by the local Ethical Committee (Comite d’ ethique en experimentation animale Charles Darwin N°5). C57BL/6J 5-week-old males were purchased from Charles River (France). Mice were housed in cages of 5, at 22LJ±LJ1LJ°C, and a 12-h light-dark cycle. Humidity levels were between 45 and 55%. Food and water were available *ad libitum.* Chronic treatment was conducted during the second half of the light phase. At 6 weeks, mice were assigned to one of the treatment groups detailed below. For sub chronic inhibition of the serotoninergic and dopaminergic system, mice were treated for 14 days with either a vehicle solution (control group), a selective 5HT-2AR antagonist (MDL100907, 2mg/kg), and a selective D2-R antagonist (Sulpiride, 25mg/kg). For sub chronic inhibition/activation of the glutamatergic system, mice were treated for 7 days with either a vehicle solution, a NMDAR antagonist (MK801, 0.3mg/kg), a mGluR5 potentiator (CDPPB, 10mg/kg). Drugs were purchased from Tocris Bioscience (CDPPB and Sulpiride) and Sigma-Aldrich (MDL 100907 and MK801). Agents were dissolved in a vehicle solution that was composed of 80% saline, 10%DMSO, 10% Cremophor and then were administered via intraperitoneal injection in mice daily at a volume equal to 10mL/kg.

### Animal tissue (brain) collection

Prior to sacrificing the mice, an intraperitoneal (i.p.) injection of pentobarbital (200mg/kg in 5% glucose) was used to fully anesthetized the animals. Brain was collected and was immediately frozen in dry ice. For RNA extraction, reverse transcription, and qPCR procedures, the protocols used were described on their respective paragraph earlier in the beginning of this section.

### Statistical Analysis

The Prism statistical analysis software (Graphpad Software Inc) was used for all statistical analysis. Examination of normality of data distribution was carried out using the Shapiro-Wilk normality test. For comparisons between two groups where the direction of change was unknown a two-tailed t-test (parametric distribution of data) or two-tailed Mann-Whitney test (non-parametric distribution of data) was used. For comparison between two groups where the direction change was already known one-tailed t-test or Mann-Whitney tests were used. For comparison of more than two groups an ordinary one-way ANOVA with post-hoc Dunnett’s multiple comparisons test (comparing all to a control group) was used for data with normal distribution, while a Kruskal Wallis ANOVA with Dunn’s multiple comparisons test (comparing all to a control group) was used. For Supplementary Fig. 2d, where comparisons between all three groups were conducted, an ordinary one-way ANOVA with Tukey’s multiple comparisons test was used. For comparison between 8 week and baseline expression data within the same patients a two-tailed paired t-test was used. For comparison of 8-week vs baseline ratios a two-tailed one-sample t-test was used (compared to mean = 1). Prism’s ROC curve calculator was used for calculating AUC and related p-values. The MedCalc statistical software was used for calculating PPV, NPV, and accuracy values.

## Results

### Blood expression levels of a brain-enriched circRNA are reliable and specific predictors of response and remission after sertraline treatment at baseline

To determine whether brain-enriched circRNAs could serve as blood biomarkers for predicting response to antidepressants, we extracted RNA from whole blood PAXgene samples from a large well-characterized antidepressant response US biomarker study, known as Establishing moderators and biosignatures of antidepressant response in clinical care (EMBARC).^31^ We then performed RNA clean-up and reverse transcription, followed by qPCR (Fig. 1a) with validated circRNA-specific primers aimed at the unique backspliced junction of a subset of previously identified brain-enriched circRNAs found to present in blood and to be linked to brain function and/or cognition (the exact selection criteria data are not shown to protect the identify of circRNA candidate biomarkers). We initially focused on the sertraline arm of the EMBARC study (Fig. 1b) and measured the expression of such candidate brain-enriched circRNA biomarkers at baseline (before treatment). Clinical response to sertraline was then determined as a 50% or more improvement in the HAMD-17 depression severity score at 8 weeks after treatment (Fig. 1b). We found that baseline blood levels of circRNAx (arbitrary circRNA identification to protect identity), a circRNA that was particularly enriched in human brain (Supplemental Fig. 1a), exhibited a significant difference in patients that responded to sertraline treatment (SERT-R) compared to patients that did not respond (SERT-NR) (Figs 1c-e). Specifically, in our discovery cohort of 49 baseline blood samples from patients treated with sertraline we found a significantly higher expression (close to 80%) in circRNAx in SERT-NR vs SERT-R (Fig. 1c). We then quantified circRNAx in another set of 42 baseline samples from the EMBARC cohort (replication cohort) and uncovered a similar upregulation (close to 70%) in SERT-NR vs SERT-R samples (Fig. 1d). Looking at the totality of the sertraline baseline samples (N = 91), we observed a clear distinction between SERT-R and SERT-NR based on baseline circRNAx blood expression (Fig. 1e; p = 0.0003 based on two-tailed Mann Whitney test). Such results were specific to this circRNA, since expression levels of circCRY2, another brain-enriched circRNA included in our study (Supplemental Fig. 1b), did not exhibit any difference between SERT-R and SERT-NR at baseline (Fig. 1f; similar results found but data not shown for a few other such circRNA candidates). Looking at various demographics such as sex, age, and race, we did not observe any significant effects on circRNAx blood expression (Supplementary Fig. 2a-d and data not shown). Interestingly, differences in baseline circRNAx levels between SERT-R and SERT-NR were also significant in the subset of patients of African American/ Asian/ Pacific Islander race (Supplementary Fig. 2c), suggesting that the predictive value of this biomarker is consistent across the race groups examined in this study.

**Fig. 1:**
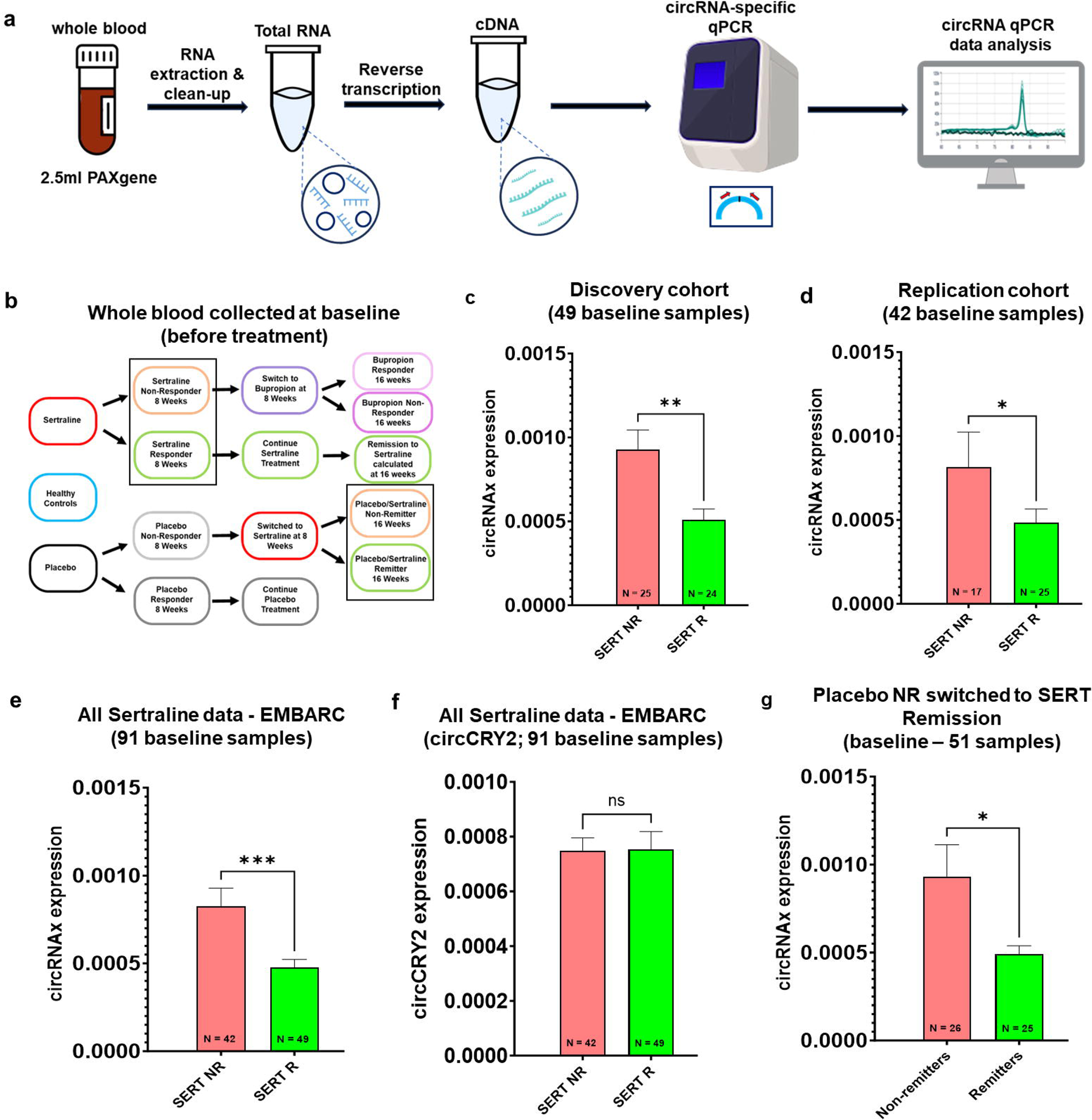
Baseline blood circRNA levels predict response and remission following sertraline treatment. Schematic of the experimental design (**a**). Human whole blood samples (PaxGene RNA IVD tubes) were obtained, subsequent RNA extraction and RNA-cleanup was performed followed by reverse transcription and circRNA-specific qPCR and quantification (see also Materials and Methods). Schematic of the EMBARC antidepressant response study (**b**). Rectangular shape indicates from arms of the study relative to the graphs included in this figure. Baseline whole blood circRNAx expression levels in SERT-R and SERT-NR in discovery (**c**) and replication (**d**) experiments with the EMBARC study. Baseline whole blood circRNAx levels in the totality of the sertraline baseline samples (**e**). Baseline whole blood expression of circCRY2 in the totality of the sertraline baseline samples (**f**). Baseline whole blood circRNAx levels in Placebo non-responder patients who were then switched to sertraline at 8 weeks and were examined for remission with sertraline treatment at 16 weeks (patients separated to Remitters and Non-Remitters to treatment). For c-e, *p < 0.05, **p < 0.01, ***p < 0.001, based on two-tailed Mann-Whitney test. For g, *p < 0.05, based on one-tailed Mann-Whitney test. Each graph is shown as Mean + SEM with the number of individual biological samples included within each graph.

To further validate that circRNAx can be a reliable biomarker for predicting response and remission with sertraline treatment, we focused on baseline samples from EMBARC MDD patients that were initially treated with Placebo, which due to lack of response were then switched to sertraline at 8weeks. Determination of remission after SERT treatment was then made at 16 weeks (HAMD17<8) (i.e. 8 weeks after initiation of SERT treatment; see also Placebo arm of the study in Fig. 1b). Our results in these sertraline-treated patients of the Placebo arm of the study indicated that blood levels of circRNAx were significantly lower at baseline in patients that achieved remission with SERT vs patients that did not (Fig. 1g), in agreement with our data in SERT-R and SERT-NR patients from the sertraline arm of the study. We conclude that circRNAx is associated with prediction of response and remission with sertraline treatment.

Focusing on the Placebo arm of the EMBARC study, we then measured circRNAx baseline levels in responders (PLA-R) and non-responders (PLA-NR) to 8 weeks of Placebo treatment using the same clinical criteria that were used for determination of response to sertraline (Fig. 2a). We observed no difference in baseline circRNAx levels in PLA-R and PLA-NR (Fig. 2b). To further test the specificity of circRNAx, we then looked at the Bupropion arm of the EMBARC study (Fig. 2a) and we separated SERT-NR patients to those that achieved remission following a subsequent 8-week treatment with the atypical antidepressant Bupropion (BUP-R) and those that did not achieve remission following Bupropion treatment after failing to respond to sertraline (BUP-NR; see also Fig. 2a). Our results suggested that baseline levels of circRNAx were not indicative of remission following Bupropion treatment after failure to respond to sertraline (Fig. 2c). We then quantified the baseline expression of circRNAx in healthy unaffected Controls and compared them with all MDD patients. Our results indicated that circRNAx was not differentially expressed between MDD patients and Controls (Fig. 2d).

**Fig. 2:**
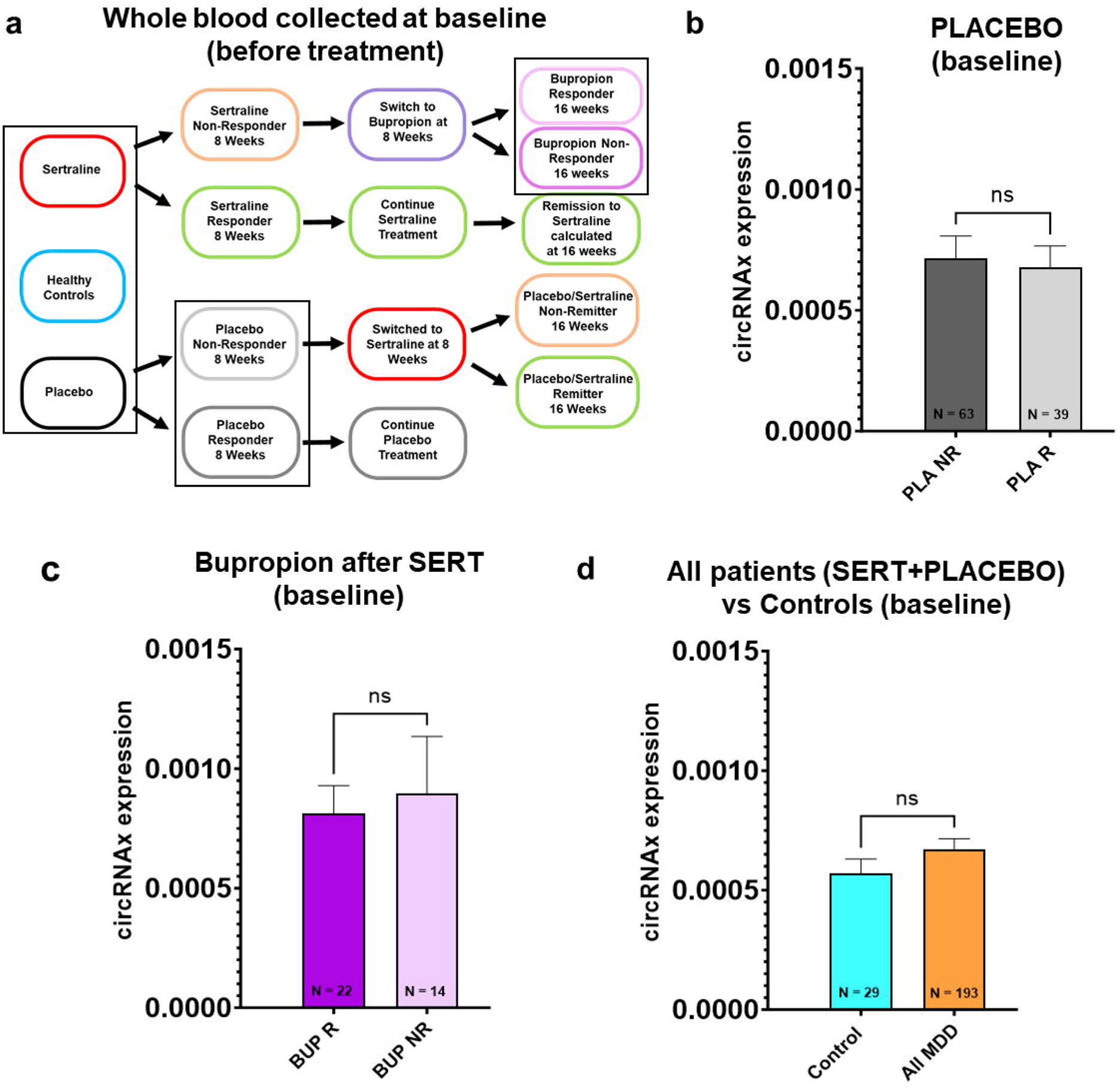
Blood circRNA levels are not associated with other treatments and are not linked to MDD diagnosis. Schematic of the EMBARC antidepressant response study (**b**). Rectangular shape indicates from arms of the study relative to the graphs included in this figure. Baseline whole blood circRNAx expression levels circRNAx in Placebo responders (PLA-R) and non-responders (PLA-NR) (**b**). Baseline whole blood circRNAx expression levels circRNAx in Bupropion responders (BUP-R) and non-responders (BUP-R) (**c**). Whole blood circRNAx baseline levels in healthy unaffected Controls and all MDD patients combined (**d**). For b-d, a two-tailed Mann-Whitney test was performed. Each graph is shown as Mean + SEM with the number of individual biological samples included within each graph.

Going back to the sertraline arm and taking advantage of the longitudinal nature of the EMBARC study sample collection, we measured the expression of circRNAx after 8 weeks of sertraline treatment and compared it to its levels at baseline within each patient. We found that levels of circRNAx displayed a significant increase after 8 weeks of sertraline treatment in the blood of SERT-R (Fig. 3a), but not SERT-NR (Fig. 3b) patients. Given that response to sertraline does not guarantee remission, we separated our SERT-R patients to remitters and non-remitters and compared the ratio of circRNAx levels at 8 weeks versus baseline for each patient for whom samples from both intervals were available. Our results indicated that SERT-R patients that achieved remission displayed a significant upregulation in circRNAx levels after 8 weeks of treatment (Fig. 3c). Importantly, no such changes were seen in SERT-R non-remitters (Fig. 3c), suggesting that circRNAx is responsive to treatment and displays a dynamic expression that can be used for disease monitoring and prediction of remission. Of note, no association was observed between changes in circRNAx levels at 8 weeks versus baseline and the possibility of remission in SERT-NR patients (Fig.3d; patients receiving Bupropion treatment after failure to respond to sertraline). Taken together, our results suggest that circRNAx is a specific, and dynamic biomarker associated with response and remission with sertraline treatment.

**Fig. 3:**
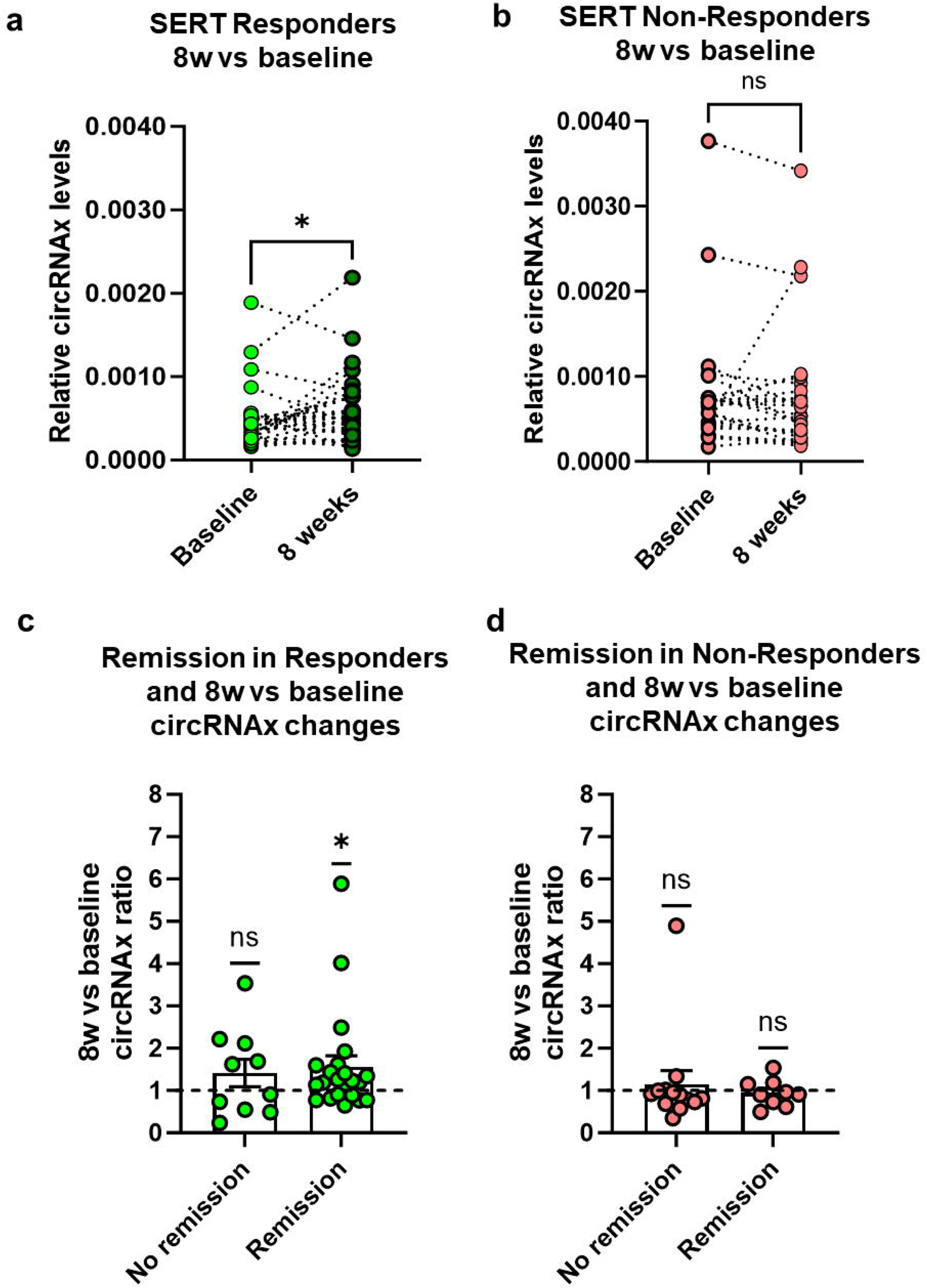
Longitudinal changes circRNAx expression levels are associated with remission with sertraline treatment. Whole blood circRNAx levels in baseline and 8 weeks of sertraline treatment in SERT-R (**a**) and SERT-NR (**b**). Ratio of 8 weeks vs baseline whole blood circRNAx levels in SERT-R patients that achieved or did not achieve remission with sertraline (**c**). Ratio of 8 weeks vs baseline whole blood circRNAx levels in SERT-NR patients that achieved or did not achieve remission with subsequent Bupropion treatment (**c**). For a-b, *p < 0.05, based on two-tailed paired t-test. For c-d, *p < 0.05, based on two-tailed one sample t-test (compared to mean ration of 1). Each graph is shown as Mean + SEM with individual biological samples values included within each graph.

### Accuracy of circRNAx blood assay for predicting response to sertraline and capacity for SSRI class prediction

To quantify the accuracy of circRNAx as a potential direct predictor of response assay for sertraline treatment, we performed a receiver operating characteristic (ROC) curve analysis on our baseline sertraline patient samples and calculated the area under the curve (AUC), as well as the positive predictive value (PPV), negative predictive value (NPV), and corresponding accuracy of circRNAx as a single analyte (Figs 4a-d). We found an AUC of 0.7153 (p = 0.0004) in the totality of the 91 baseline samples examined (Fig. 4a) by utilizing a single cut-off for discriminating between SERT-R and SERT-NR. Given the known heterogeneity of MDD, we hypothesized that confounding factors related to misdiagnosis or different subtypes of MDD could be affecting the performance of our assay. Given that a small subset of patients initially diagnosed with MDD end up being later diagnosed with Bipolar Disorder,^33^ we looked for potential clinical indicators of Bipolar Disorder susceptibility within our cohort (analysis was limited to female patients since they represented close to 70% of our sertraline cohort). Given the strong hereditary nature of Bipolar Disorder,^34^ we focused on the presence or absence of family history of mania. We also included in our analysis the HAMD-31 clinical question that interrogates about the potential circadian nature of depression symptoms, which itself is indirectly associated with the presence of atypical depression (HAMD-31 score of 3 = onset of depression symptoms late during the day).^35^ Interestingly, we found that excluding patients that had a positive family history of mania (10% of our sertraline cohort) and patients that had a score of 3 in the HAMD-31 question (11% of our cohort) resulted in a significantly improved performance for our assay (Figs. 4b-d; AUC = 0.8083, p-value < 0.0001). Specifically, we found that the PPV increased to 80.95% from 70.18% and the NPV increased to 77.42% from 73.53%, for a total accuracy of 79.45% (Fig. 4c). Notably, including any of the two clinical questions alone, increased the accuracy of the assay to 75-76% (Fig. 4c). Even though levels of circRNAx were not affected by sex, themselves (see also Supplementary Fig. 2b), we noticed that the predictive value of our assay was much stronger in male than female patients without the inclusion of the above clinical questions (Fig. 4d). Importantly, the inclusion of the family history of mania and HAMD-31 score questions, managed to significantly elevate the accuracy of our assay for female patients (Fig. 4d). These data suggest, that circRNAx can be used as a single biomarker with acceptable accuracy for predicting response to sertraline treatment.

**Fig. 4:**
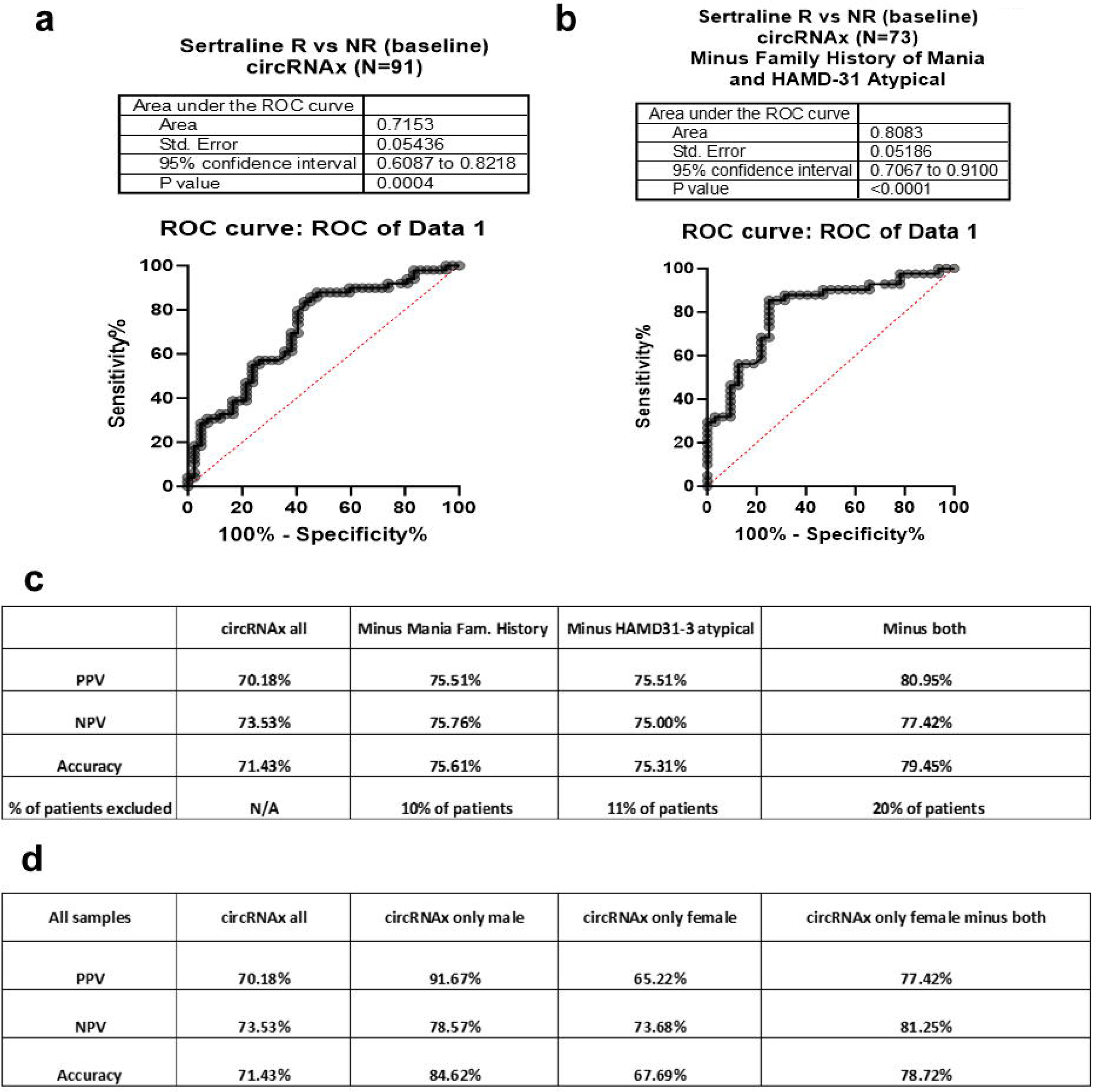
Accuracy of circRNAx blood assay for predicting response to sertraline. ROC analysis curve for circRNAx baseline levels between SERT-R and SERT-NR (**a**). The AUC (Area Under the Curve) along with the table of statistics is included ROC analysis curve for circRNAx baseline levels between SERT-R and SERT-NR after eliminating female patients with positive family history of mania and a HAMD-31 score of 3 (**b**) (onset of depression symptoms late during the day). Table showing the PPV, NPV, and accuracy of the circRNAx assay in all patients (circRNAx all), after elimination of female patients with positive family history of mania, after elimination of subjects with a HAMD-31 score of 3, or after elimination of patients with either a positive family history of mania or a HAMD-31 score of 3 (**c**). Table showing the PPV, NPV, and accuracy of the circRNAx assay in all patients regardless of sex (circRNAx all), only in male patients (circRNAx only male), only in female patients (circRNAx only female), and in female patients after elimination of patients with positive family history of mania, after elimination of subjects with a HAMD-31 score of 3, or after elimination of patients with either a positive family history of mania or a HAMD-31 score of 3 (circRNAx only female minus both) (**d**).

Although a reliable and accurate biological biomarker for predicting response to sertraline could result in significant benefits to the existing clinical workflow, a prediction of response to the whole class of SSRIs might provide physicians with additional flexibility for effective personalized treatment selection. To that end, and in order to further validate the predictive value of circRNAx in additional sample types, we utilized total RNA extracted from leukocytes collected from venous blood of patients with MDD treated with various classes of antidepressants as part of a naturalistic clinical study (Supplementary Fig. 3a).^32^ Focusing on MDD patients who were prescribed various SSRIs and for which remission was calculated at 30 weeks after treatment, we found that circRNAx levels were significantly higher in non-remitters vs remitters (Supplementary Fig. 3b), and that despite the smaller size of this cohort, appeared to have a similar predictive effect (Supplementary Fig. 2c; AUC=0.8750, p = 0.0550). Importantly, focusing on another subset of patients who were treated with other classes of antidepressants (SNRIs, TCAs, MAOIs), we observed no significant effects on circRNAx levels and remission following treatment (Supplementary Fig. 3d). Furthermore, levels of circRNAx in leukocytes from patients with MDD and Bipolar Disorder were not significantly different from unaffected healthy Controls (Supplementary Fig. 3e). We conclude that circRNAx levels in leukocytes can specifically predict remission with the SSRI class of antidepressants.

### Levels of circRNAx in mouse brain are regulated by serotonin receptor 5-HT2A activity

Even though a reliable molecular biomarker does not need to be necessarily associated with known disease-related molecular pathways, we decided to further investigate the potential mechanisms that could underlie the biogenesis of circRNAx within the brain. To that end, we took advantage of the fact that circRNAx is partially conserved between human and mouse and significantly enriched in mouse brain and neuronal cultures. We then treated mice with specific inhibitors targeting different neuronal receptor s (glutamatergic NMDA receptor (MK801), D2 dopaminergic receptors (sulpiride), and 5-HT2A receptors (MDL100907). These receptors are important for brain functions and cognition and relevant to the pathophysiology of depression (Fig. 5a).^36–40^ We also included treatment with the mGluR5 positive allosteric modulator, CDPPB. We found that serotonin 5-HT2A receptor but not D2 receptor antagonism was able to significantly downregulate circRNAx in all three of the brain regions examined in this study (frontal cortex, nucleus accumbens, and putamen; Figs. 5b-c and Supplementary Fig. 4a). Furthermore, no effects were observed in mouse brain circRNAno1 levels following either NMDA receptor antagonism or mGLUR5 receptor activation (Figs. 5d-e). Of note, focusing on another brain-enriched circRNA (circTulp4)^13^, we found no significant changes as a result of 5-HT2A receptor blockade, but an upregulation following D2 receptor antagonism in two for the three brain regions examined (Supplementary Figs. 4b-d). On the other hand, a significant downregulation in circTulp4 expression was observed in the nucleus accumbens of mice treated with MK801 (Supplementary Figs. 4e-f), suggesting that different brain-specific circRNAs are sensitive to diverse types of neuronal receptors. Our results suggest that circRNAx expression in the mouse brain is regulated by serotonin receptor activity but is not significantly affected by glutamatergic NMDA/mGLUR5 and dopaminergic D2 receptor function.

**Fig. 5:**
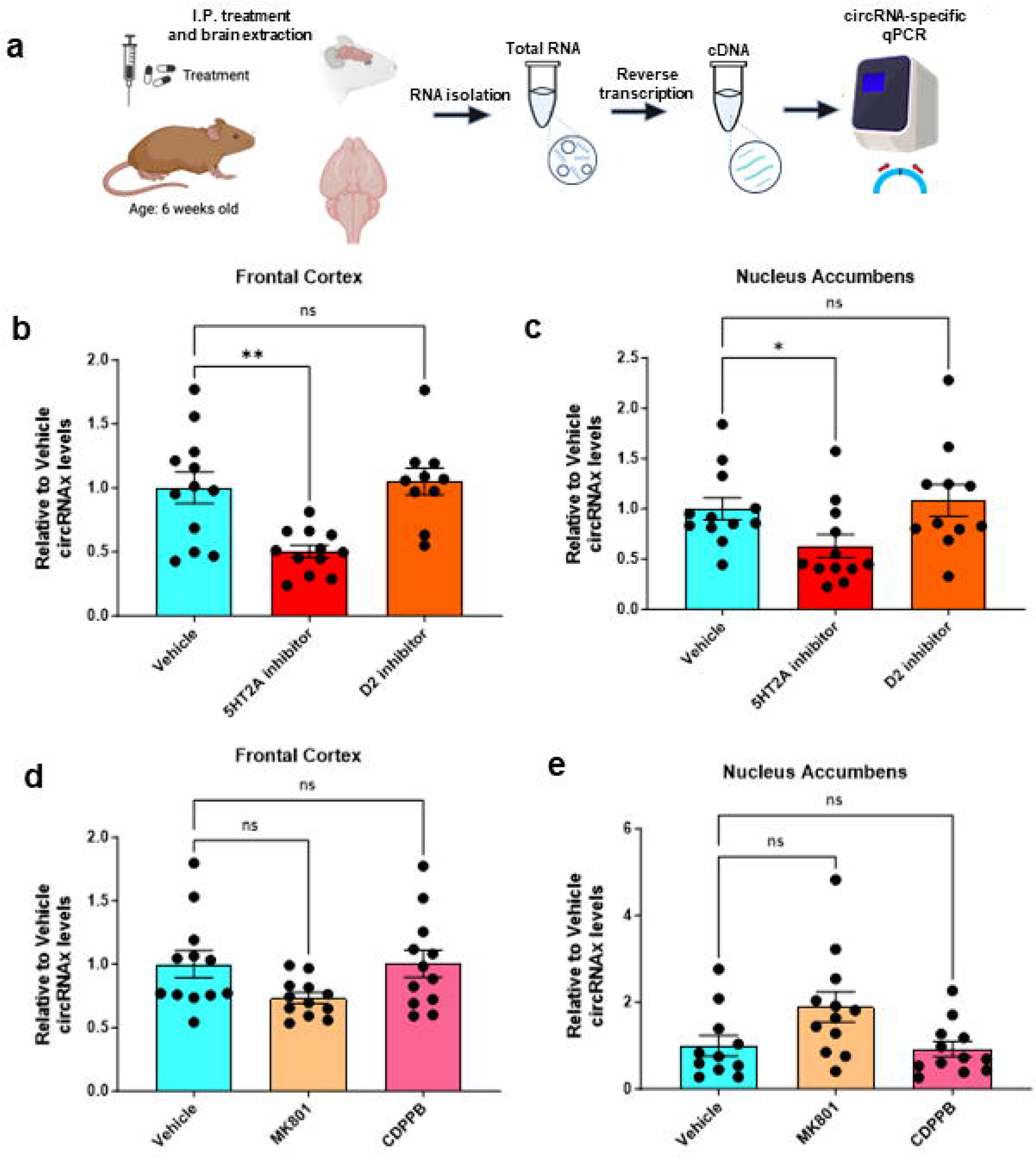
circRNAx levels in mouse brain are inhibited following serotonin 5-HT2A receptor antagonism. Experimental design scheme (**a**). WT 5-week-old mice were treated with various pharmacological agents that aim the blockade of neuronal brain receptors. Mice were euthanized and different brain regions were dissected. Brain tissue from brain regions of interest was subjected to RNA extraction, reverse transcription, and circRNA qPCR. Mouse brain circRNAx levels after treatment with a pure 5T2AR antagonist MDL100907 and the pure D2R antagonist Sulpiride in mouse frontal cortex (**b**) and nucleus accumbens (**c**). Mouse brain circRNAx levels after treatment with MK801, a selective NMDAR antagonist and CDPPB, an mGluR5 potent agonist in mouse frontal cortex (**d**) and nucleus accumbens (**e**). *p < 0.05, **p < 0.01, based on an ordinary one-way ANOVA with post-hoc Dunnett’s multiple comparisons test (b) or Kruskal Wallis ANOVA with Dunn’s multiple comparisons test (c). Each graph is shown as Mean + SEM with individual biological samples values included within each graph.

### Neuronal expression of CircRNAx is regulated by BDNF receptor activity and involves the ERK/CREB signaling pathway

To further dissect the molecular pathways that could underlie the control of circRNAx expression within neurons, we utilized mouse cortical neuronal cultures and human neuroblastoma cell lines that were treated with different chemical inhibitors of molecular pathways associated with neuronal activity (Fig. 5a-f)^41^. We initially tested the effects of BDNF receptor (trkB) inhibition in human neuroblastoma SH-SY5Y cell lines, so as to determine if circRNAx could be downstream of neurotrophic cellular signaling (Fig. 6a). Our results showed that TtrkB inhibition can significantly downregulate circRNAx expression in human neuroblastoma cells (Fig. 6b). This effect was specific, since levels of circTulp4 were not affected by TrkB inhibition (Fig. 6c). Importantly, repeating TrkB inhibitor treatment in mouse cortical neurons lead to a significant downregulation of circRNAx expression (Fig. 6d-e). To further dissect the molecular pathways that could be associated with the control of neuronal circRNAx expression, we treated cortical neurons with CBP/CREB and ERK inhibitors. We found that circRNAx was significantly downregulated by both treatments (Fig. 6e), suggesting that ERK/CREB signaling is upstream of circRNAx synthesis. Additional treatments with PKA and PKC inhibitors showed no effects on circRNA expression in mouse cortical neurons (Supplementary Fig. 5a-b). Taking together all in vitro and in vivo experiments we can surmise that circRNAx expression in mouse neurons is activated via 5-HT2A-and BDNF/TrkB-mediated activation of ERK/CREB signaling (Supplementary Fig. 6).

**Fig. 6:**
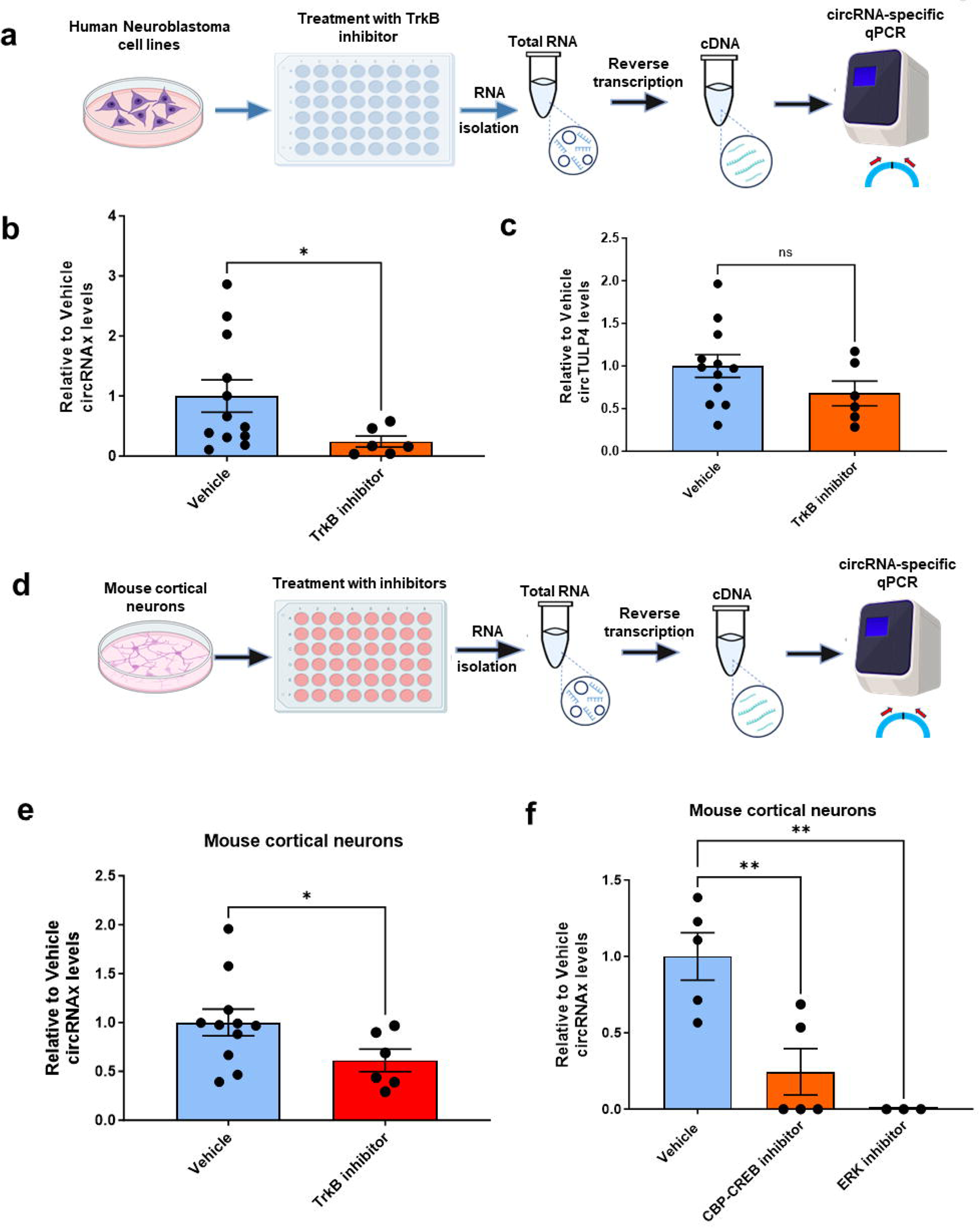
Neuronal circRNAx levels are influenced by BDNF receptor activity and ERK/CREB signaling. Schematic of the experimental design for the human neuroblastoma cell line (**a**). SHSY5 cells were cultured, plated in a well plate, and treated with various pharmacological inhibitors. At 24hrs after treatment, cells were harvested, RNA was extracted, and reverse transcription and circRNA-specific PCR was performed. CircRNAx (**b**) and circTulp4 (**c**) levels in human neuroblastoma SHSY5 cells after treatment with TrkB receptor antagonist. Experimental Design for mouse primary mouse cortical neuron experiments (**d**). CircRNAx levels in primary mouse cortical neurons after treatment with TrkB receptor antagonist (**e**). CircRNAx levels in primary mouse cortical neurons after treatment with CREB-CBP interaction inhibitor and a potent and selective ERK inhibitor (**f**). For (a) and (e): *p < 0.05 based on two-tailed (a) and one-tailed (e) Mann-Whitney test. For (f): **p < 0.01, based on an ordinary one-way ANOVA with post-hoc Dunnett’s multiple comparisons test. Each graph is shown as Mean + SEM with individual biological samples values included within each graph.

## Discussion

Despite the progressive evolution in our understanding of the biological mechanisms associated with the pathogenesis of MDD and the further elucidation of molecular pathways with relevance to antidepressant response, there has been limited progress in the discovery of reliable and robust biological biomarkers with clinical relevance for MDD diagnosis and treatment. Here we provide data suggesting that baseline blood levels of a brain-enriched circRNA (circRNAx) associated with serotonin and BDNF receptor activity, can significantly predict response and remission following treatment with SSRI antidepressants. We further highlight the accuracy of a single analyte circRNA assay and demonstrate its specificity for this class of antidepressants. We provide additional mechanistic understandings of the molecular pathways that could regulate the expression of this circRNA in the CNS. Taken together, our data introduce a novel biological biomarker for predicting response to antidepressant treatment that is significantly associated with molecular cascades of relevance to antidepressant response within the brain.

Our data clearly demonstrate a robust enrichment in circRNAx levels in human brain vs other human organs (see also Supplementary Fig. 1), which agrees with previous studies, whose circRNA sequencing data include circRNAx among the circRNAs found to be enriched in human brain.^11,13^ Furthermore, circRNAx is among the most readily-detected circRNAs in human cerebrospinal fluid, suggesting that it can cross the BBB.^12,42^ Additional insights into the mechanisms of circRNAx secretion and trafficking are studies suggesting a significant enrichment of this circRNA in exosomes,^43^ which themselves are known to be able to cross the BBB.^44^ The combination of particularly high levels of this circRNA in the human brain, its capacity to cross the BBB, and the very low expression of this circRNA in other human tissues, makes highly likely that levels of circRNAx in the blood are mostly derived from the brain. The fact that circRNAx levels in the blood appear to respond to sertraline treatment and our findings that serotonin and BDNF receptor antagonism can reduce mouse brain circRNAx levels further support this scenario.

Our EMBARC clinical study data suggest that whole blood levels of circRNAx are lower in SERT-R vs SERT-NR and that sertraline treatment leads to a significant upregulation in blood circRNAx levels only in SERT-R patients; an effect that is also associated with possibility for remission with sertraline. Of note, levels in unaffected Controls are similar to SERT-R levels, suggesting that physiological baseline levels of blood circRNAx are needed for response to sertraline and subsequent increase in such levels is required for remission with treatment. On the other hand, non-responders to sertraline show saturated levels in circRNAx expression that remain unchanged after treatment. Furthermore, the predictive value of circRNAx was limited to sertraline (EMBARC whole blood study) or the SSRI class of antidepressants (leukocyte clinical study) and not associated with response to placebo or other classes of antidepressants. Based on our animal and neuronal culture mechanistic data, circRNAx is downstream of serotonin and BDNF receptor signaling, but is not substantially influenced by dopamine or glutamatergic neurotransmission. Given that serotonin and BDNF signaling appear to be more closely linked to known effects of SSRI treatment within the brain,^36,37^ such findings suggest that circRNAx could be specifically linked to mechanisms of antidepressant response with relevance to the SSRI class of antidepressants. It is, thus, tempting, to hypothesize that the capacity to dynamically regulate circRNAx levels in brain and blood over a significant threshold after treatment with SSRIs could provide a potential explanation for our findings. However, additional studies are required to further dissect the molecular mechanisms that could underlie the control of brain circRNAx expression and secretion into blood.

Our animal and neuronal culture studies identified the 5-HT2A and BDNF/TrkB receptors as upstream regulators of circRNAx, as inhibition of both such receptors resulted in reduced circRNAx expression. We also concluded after testing a number of intracellular components of neuronal gene expression that the ERK/CREB molecular pathways were important downstream regulators for neuronal circRNAx expression. Given that both 5-HT2A and TrkB receptor activation can result in ERK/CREB phosphorylation and activation, such downstream signaling cascades are consistent with our neuronal receptor data (see also Supplementary Fig. 6). Interestingly, other serotonin receptors not tested in our study, such as 5-HT2B, 5-HT2C, 5-HT3, and 5-HT6, could also activate the ERK/CREB signaling.^37^ It is, thus, possible that such receptors could be also of relevance for circRNAx expression in the brain. However, given that other serotonin receptors, such as 5-HT1 and 5-HT5, can have opposing effects in such intracellular neuronal cascades,^37^ we would expect that activation of such receptors would likely result in opposing effects in circRNAx expression. It is, thus, tempting to hypothesize, that circRNAx expression in the brain and blood could be specifically controlled via the synergistic activation of specific serotonin and BDNF receptors. In such a hypothetical scenario, circRNAx levels in brain and brain-derived circRNAx levels in the blood could be predictive of the capacity of an individual patient to specifically activate both such receptors following SSRI antidepressant treatment.

Given the significant brain-enrichment of circRNAx, it is possible that disorders that significantly impact brain tissue or notably disrupt the BBB, such as traumatic brain injury or stroke, could confound blood circRNAx measurements. Since the clinical samples used in our study did not allow for such significant comorbidities, we are unable to fully characterize such effects, which is a limitation of our study. Lastly, well-characterized MDD clinical studies, such as EMBARC, are designed to include strict inclusion and exclusion criteria, thus not allowing for the presence of other psychiatric comorbidities that very often accompany MDD. On a similar note, our results identified that the accuracy of our circRNAx assay is significantly improved if we exclude patients with family history of mania or atypical HAMD-31 circadian rhythmicity in MDD symptoms. Such findings suggest that reducing misdiagnosis between MDD and of other psychiatric disorders with depression symptomatology, such as Bipolar Disorder, could further increase the clinical efficacy of a molecular biomarker specifically predictive of SSRI response in MDD patients.

Previous studies have uncovered significant effects on brain gene expression and function following specific manipulations in circRNAx and its downstream molecular targets (references not included to protect identity of circRNA). It is, thus, possible, that circRNAx could be eventually proven to be an important novel molecular component of the pathophysiology of antidepressant response and future potential therapeutic target for depression. Furthermore, our data showing longitudinal changes in circRNAx levels in responders to sertraline being linked to long term remission, suggest the opportunity for this and other circRNAs to serve as biomarkers for long-term depression monitoring. Future work is, thus, needed to further harness the power of circRNAs as novel biomarkers for the better diagnosis and treatment of psychiatric and neurological disorders.

## Acknowledgements

We would like to thank Dr. Roy Perlis, Dr. Michael Ackermann, Dr. Nikolaos Daskalakis, and Dr. Carlos Cruchaga for their valuable feedback on our manuscript.

## References

1. Goodwin, R. D., et al. Trends in U.S. Depression Prevalence From 2015 to 2020: The Widening Treatment Gap. Am. J. Prev. Med. 63, 726–733 (2022).

2. Mental health in the United States | The COVID States Project. https://www.covidstates.org/reports/mental-health-in-the-united-states.

3. 3. Inc, G. U.S. Depression Rates Reach New Highs. Gallup.com https://news.gallup.com/poll/505745/depression-rates-reach-new-highs.aspx (2023).

4. Gaynes, B. N. et al. What did STAR*D teach us? Results from a large-scale, practical, clinical trial for patients with depression. Psychiatr. Serv. Wash. DC 60, 1439–1445 (2009).

5. Cipriani, A. et al. Comparative efficacy and acceptability of 21 antidepressant drugs for the acute treatment of adults with major depressive disorder: a systematic review and network meta-analysis. Lancet Lond. Engl. 391, 1357–1366 (2018).

6. Frye, M. A. & Nemeroff, C. B. Pharmacogenomic testing for antidepressant treatment selection: lessons learned and roadmap forward. Neuropsychopharmacology 49, 282– 284 (2024).

7. Salzman, J., Chen, R. E., Olsen, M. N., Wang, P. L. & Brown, P. O. Cell-type specific features of circular RNA expression. PLoS Genet. 9, e1003777 (2013).

8. Memczak, S. et al. Circular RNAs are a large class of animal RNAs with regulatory potency. Nature 495, 333–338 (2013).

9. Jeck, W. R. et al. Circular RNAs are abundant, conserved, and associated with ALU repeats. RNA N. Y. N 19, 141–157 (2013).

10. Zhang, Y. et al. The Biogenesis of Nascent Circular RNAs. Cell Rep. 15, 611–624 (2016).

11. You, X. et al. Neural circular RNAs are derived from synaptic genes and regulated by development and plasticity. Nat. Neurosci. 18, 603–610 (2015).

12. Wang, Z. et al. Clinical utility of cerebrospinal fluid-derived circular RNAs in lung adenocarcinoma patients with brain metastases. J. Transl. Med. 20, 74 (2022).

13. Rybak-Wolf, A. et al. Circular RNAs in the Mammalian Brain Are Highly Abundant, Conserved, and Dynamically Expressed. Mol. Cell 58, 870–885 (2015).

14. Wen, G. & Gu, W. Circular RNAs in peripheral blood mononuclear cells are more stable than linear RNAs upon sample processing delay. J. Cell. Mol. Med. 26, 5021–5032 (2022).

15. Memczak, S., Papavasileiou, P., Peters, O. & Rajewsky, N. Identification and Characterization of Circular RNAs As a New Class of Putative Biomarkers in Human Blood. PloS One 10, e0141214 (2015).

16. Liu, C.-X. & Chen, L.-L. Circular RNAs: Characterization, cellular roles, and applications. Cell 185, 2016–2034 (2022).

17. Hafez, A. K. et al. A bidirectional competitive interaction between circHomer1 and Homer1b within the orbitofrontal cortex regulates reversal learning. Cell Rep. 38, 110282 (2022).

18. Seeler, S. et al. A Circular RNA Expressed from the FAT3 Locus Regulates Neural Development. Mol. Neurobiol. 60, 3239–3260 (2023).

19. Suenkel, C., Cavalli, D., Massalini, S., Calegari, F. & Rajewsky, N. A Highly Conserved Circular RNA Is Required to Keep Neural Cells in a Progenitor State in the Mammalian Brain. Cell Rep. 30, 2170–2179.e5 (2020).

20. Zimmerman, A. J. et al. A psychiatric disease-related circular RNA controls synaptic gene expression and cognition. Mol. Psychiatry 25, 2712–2727 (2020).

21. Dube, U. et al. An atlas of cortical circular RNA expression in Alzheimer disease brains demonstrates clinical and pathological associations. Nat. Neurosci. 22, 1903–1912 (2019).

22. Hanan, M., Soreq, H. & Kadener, S. CircRNAs in the brain. RNA Biol. 14, 1028–1034 (2017).

23. Piwecka, M. et al. Loss of a mammalian circular RNA locus causes miRNA deregulation and affects brain function. Science 357, eaam8526 (2017).

24. Yoon, G. et al. Obesity-linked circular RNA circTshz2-2 regulates the neuronal cell cycle and spatial memory in the brain. Mol. Psychiatry 26, 6350–6364 (2021).

25. Song, R. et al. Plasma Circular RNA DYM Related to Major Depressive Disorder and Rapid Antidepressant Effect Treated by Visual Cortical Repetitive Transcranial Magnetic Stimulation. J. Affect. Disord. 274, 486–493 (2020).

26. Malhotra, S., Miras, M. C. M., Pappolla, A., Montalban, X. & Comabella, M. Liquid Biopsy in Neurological Diseases. Cells 12, 1911 (2023).

27. Ravanidis, S. et al. Differentially Expressed Circular RNAs in Peripheral Blood Mononuclear Cells of Patients with Parkinson’s Disease. Mov. Disord. Off. J. Mov. Disord. Soc. 36, 1170–1179 (2021).

28. Zhou, M., Li, S. & Huang, C. Physiological and pathological functions of circular RNAs in the nervous system. Neural Regen. Res. 19, 342–349 (2024).

29. Verduci, L., Tarcitano, E., Strano, S., Yarden, Y. & Blandino, G. CircRNAs: role in human diseases and potential use as biomarkers. Cell Death Dis. 12, 468 (2021).

30. Ma, Y., Liu, Y. & Jiang, Z. CircRNAs: A new perspective of biomarkers in the nervous system. Biomed. Pharmacother. Biomedecine Pharmacother. 128, 110251 (2020).

31. Trivedi, M. H. et al. Establishing moderators and biosignatures of antidepressant response in clinical care (EMBARC): Rationale and design. J. Psychiatr. Res. 78, 11–23 (2016).

32. Apazoglou, K. et al. Antidepressive effects of targeting ELK-1 signal transduction. Nat. Med. 24, 591–597 (2018).

33. Kessing, L. V., Willer, I., Andersen, P. K. & Bukh, J. D. Rate and predictors of conversion from unipolar to bipolar disorder: A systematic review and meta-analysis. Bipolar Disord. 19, 324–335 (2017).

34. Barnett, J. H. & Smoller, J. W. The Genetics of Bipolar Disorder. Neuroscience 164, 331–343 (2009).

35. Singh, T. & Williams, K. Atypical depression. Psychiatry Edgmont Pa Townsh. 3, 33–39 (2006).

36. Martinowich, K. & Lu, B. Interaction between BDNF and serotonin: role in mood disorders. Neuropsychopharmacol. Off. Publ. Am. Coll. Neuropsychopharmacol. 33, 73– 83 (2008).

37. Zięba, A., Stępnicki, P., Matosiuk, D. & Kaczor, A. A. Overcoming Depression with 5-HT2A Receptor Ligands. Int. J. Mol. Sci. 23, 10 (2021).

38. Dunlop, B. W. & Nemeroff, C. B. The role of dopamine in the pathophysiology of depression. Arch. Gen. Psychiatry 64, 327–337 (2007).

39. Lv, S., Yao, K., Zhang, Y. & Zhu, S. NMDA receptors as therapeutic targets for depression treatment: Evidence from clinical to basic research. Neuropharmacology 225, 109378 (2023).

40. Potter, L. E., Zanos, P. & Gould, T. D. Antidepressant Effects and Mechanisms of Group II mGlu Receptor-Specific Negative Allosteric Modulators. Neuron 105, 1–3 (2020).

41. Flavell, S. W. & Greenberg, M. E. Signaling mechanisms linking neuronal activity to gene expression and plasticity of the nervous system. Annu. Rev. Neurosci. 31, 563– 590 (2008).

42. Lee, B. et al. Medulloblastoma cerebrospinal fluid reveals metabolites and lipids indicative of hypoxia and cancer-specific RNAs. Acta Neuropathol. Commun. 10, 25 (2022).

43. Li, Y. et al. Circular RNA is enriched and stable in exosomes: a promising biomarker for cancer diagnosis. Cell Res. 25, 981–984 (2015).

44. Ramos-Zaldívar, H. M. et al. Extracellular vesicles through the blood-brain barrier: a review. Fluids Barriers CNS 19, 60 (2022).

