## Supplementary Figures 1-6 for "A brain-enriched circRNA blood biomarker can predict response to SSRI antidepressants"

**Contents:**

**Pages 2-7: Supplementary Figures 1-6**

**Pages 7-9: Supplementary Figure legends**

**Supplementary Figures**

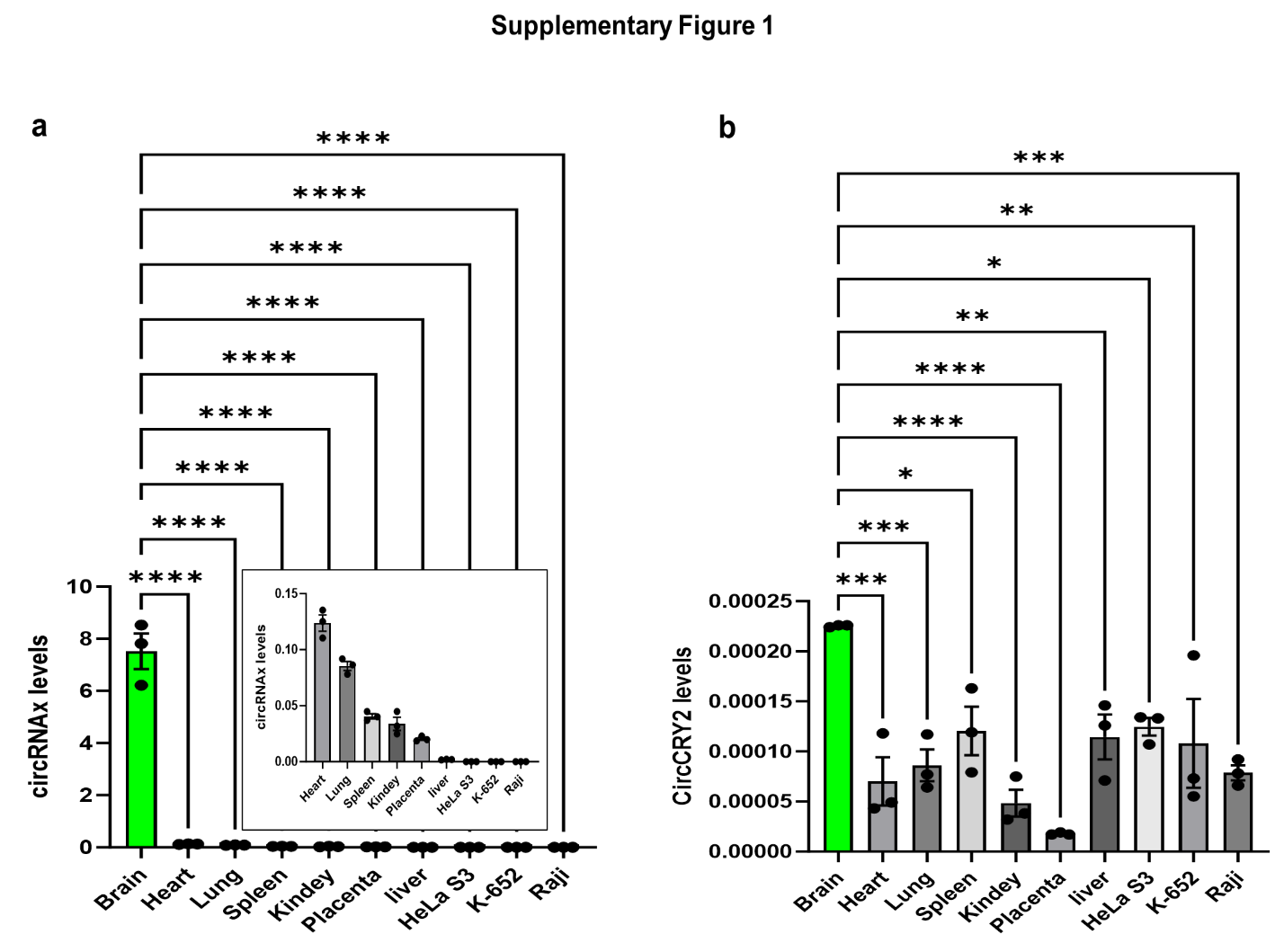

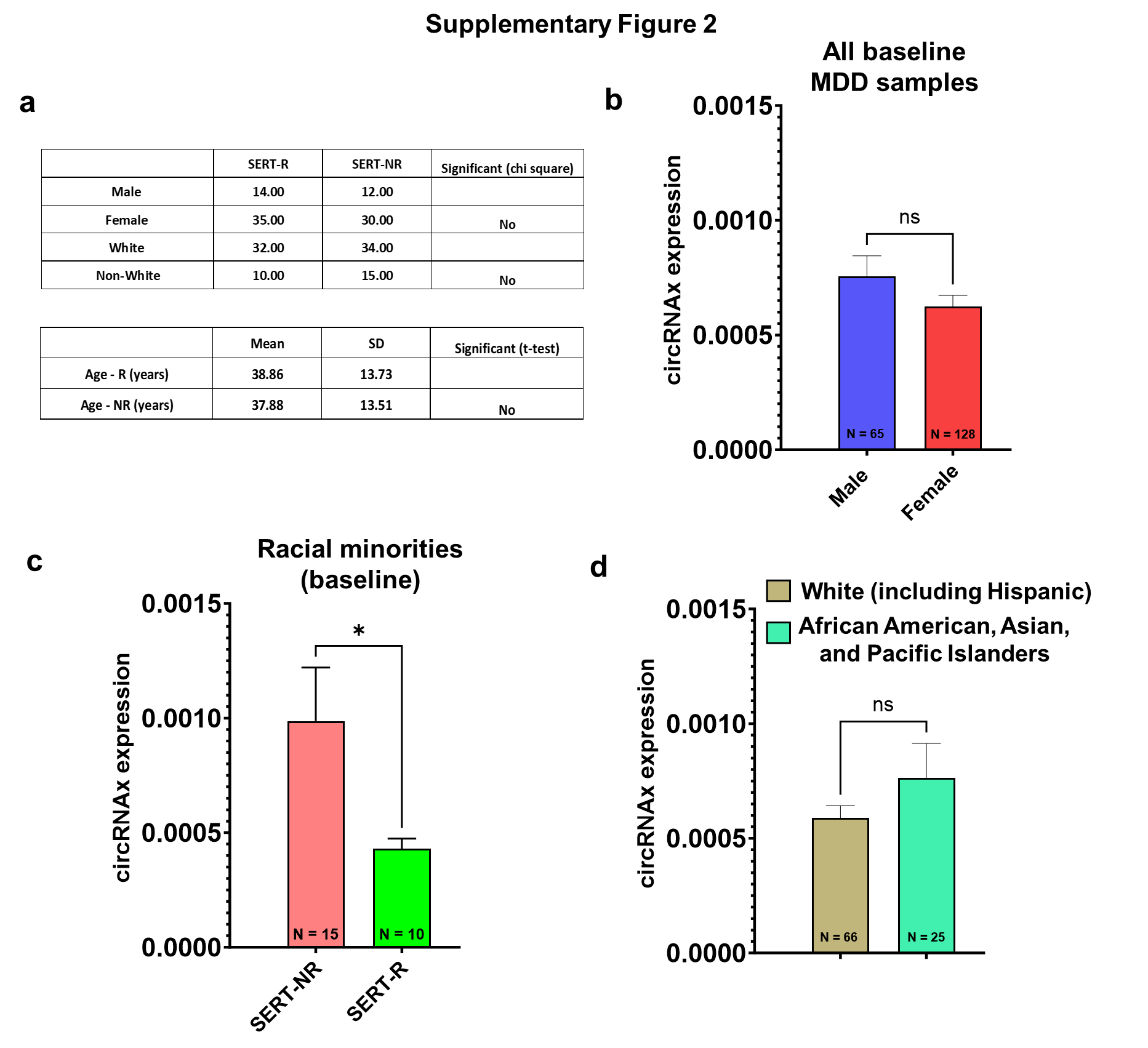

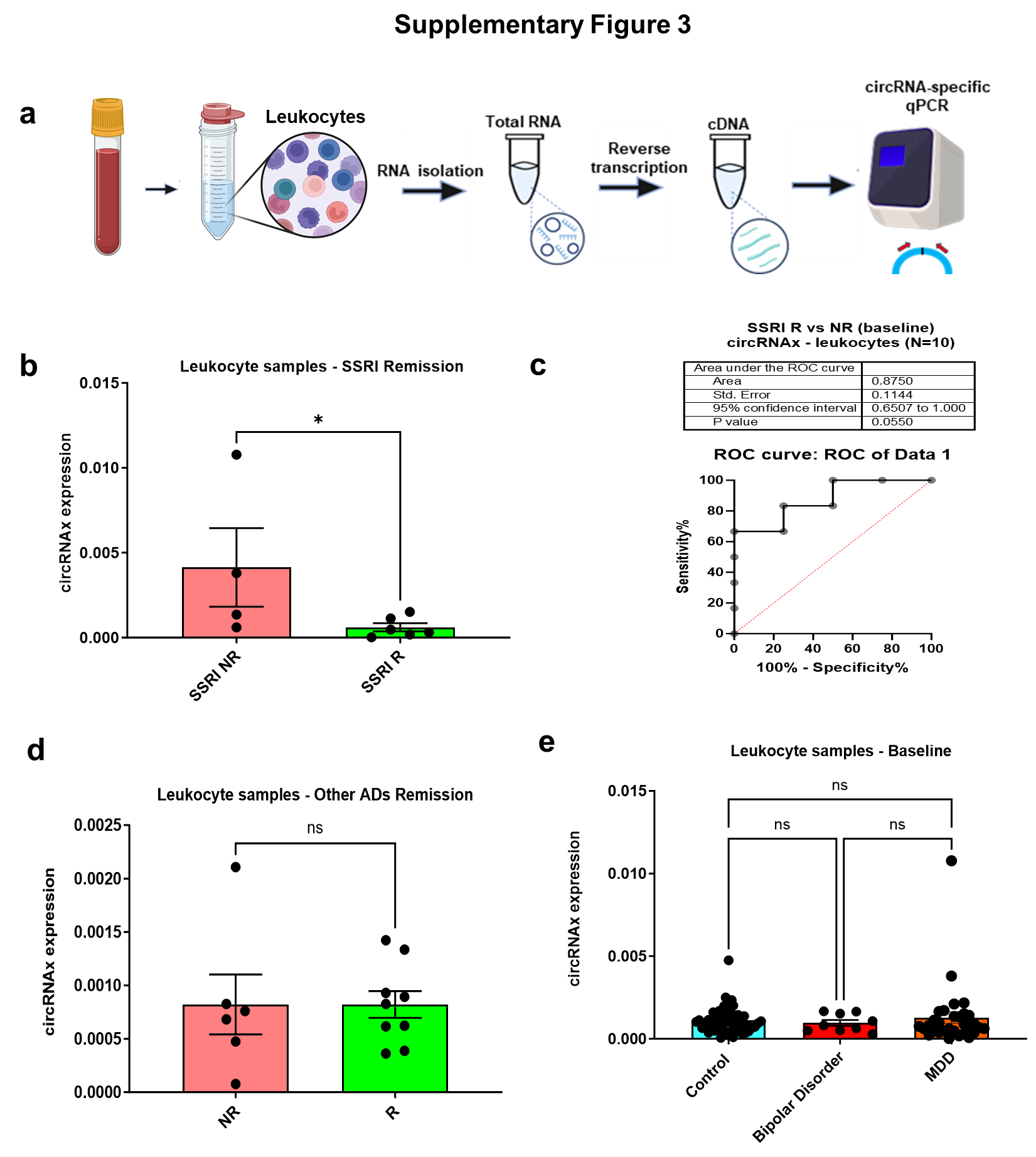

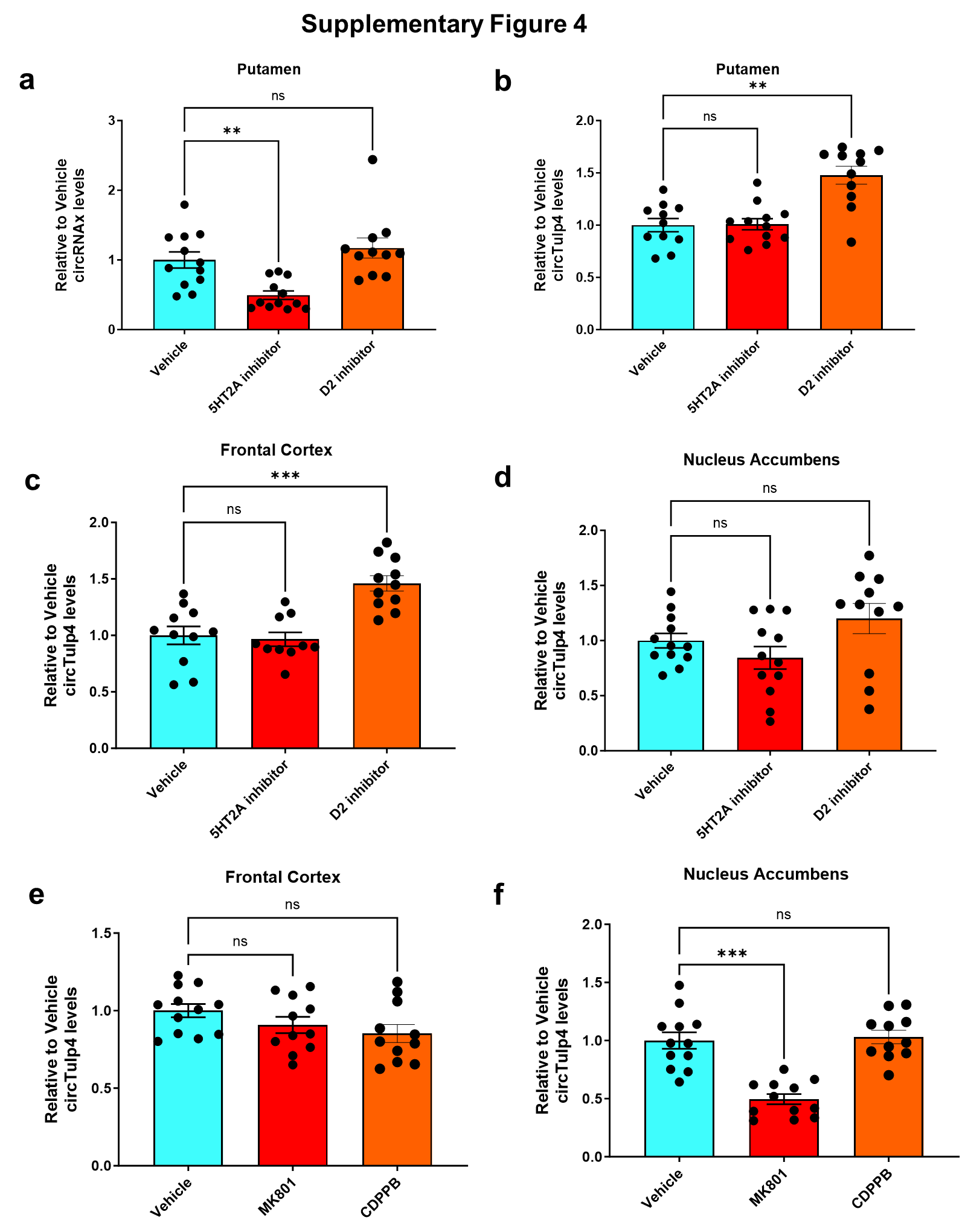

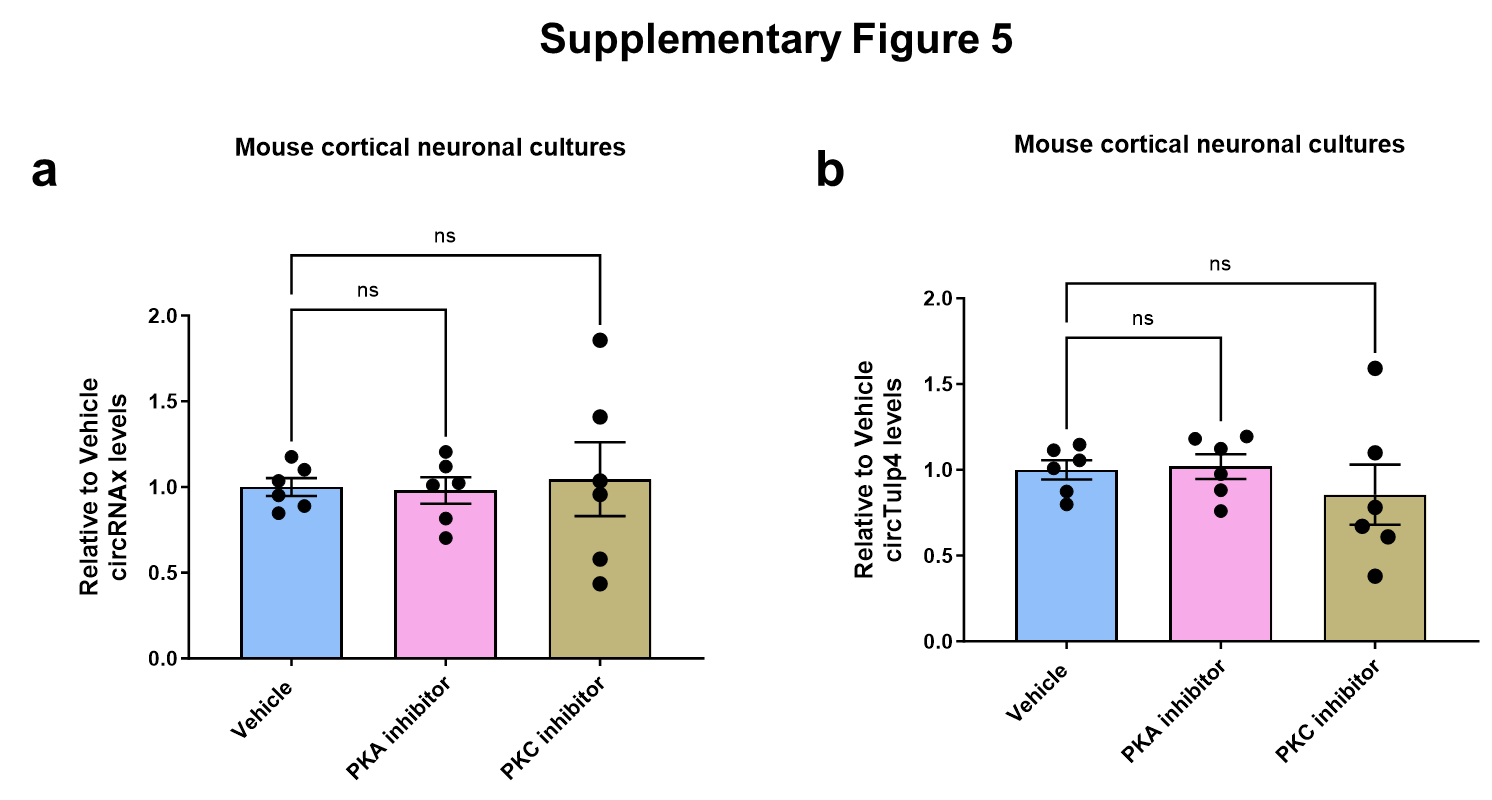

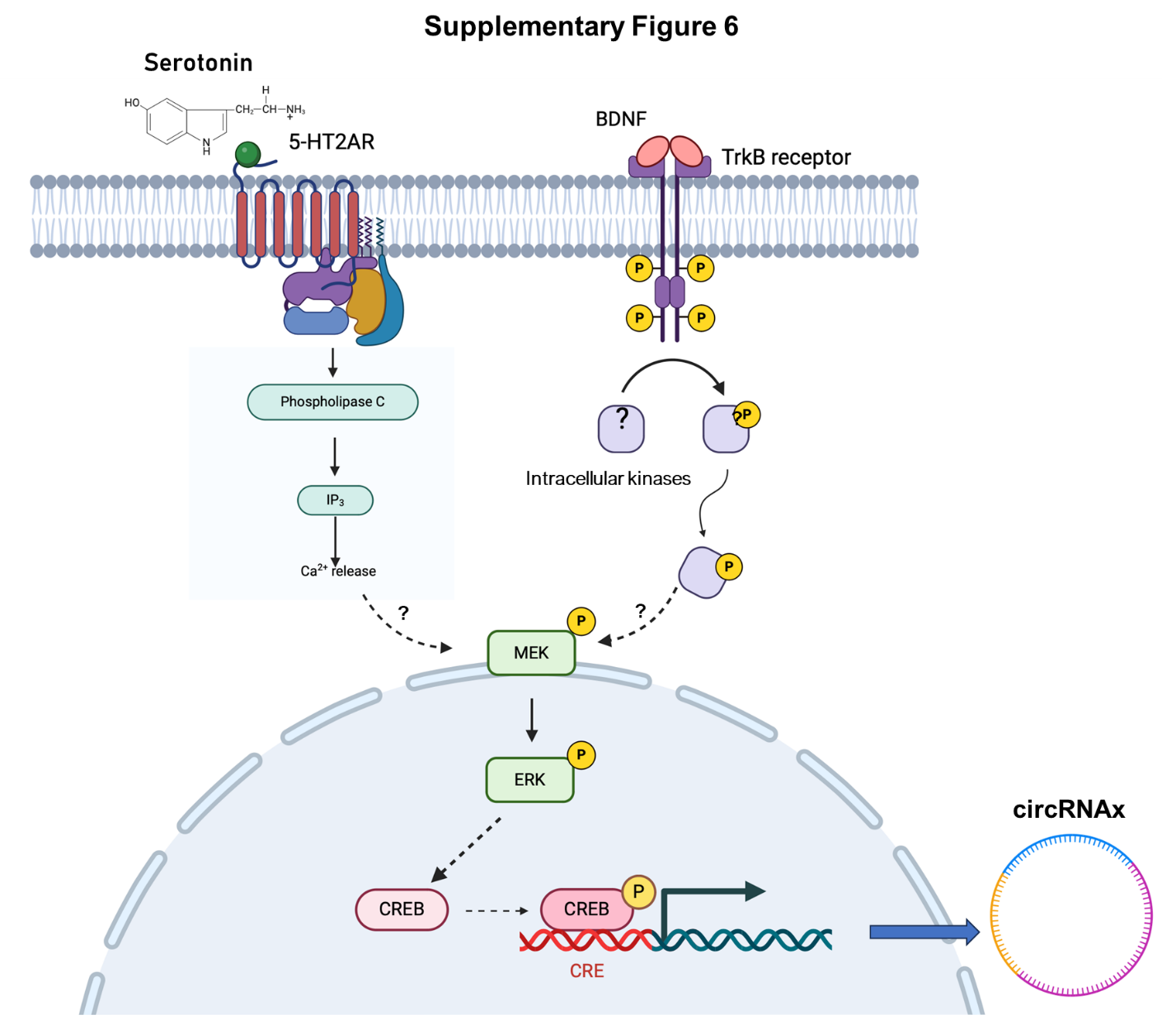

**Supplementary Figure Legends:**

**Supplementary Figure 1: Effects of demographics on circRNAx expression**

Tables showing the number of male and female patients and white and non-white patients in the SERT-R and SERT-NR groups (**a**- above), as well as the mean and SD for age and weight (**a**- below). Baseline circRNAx levels in male and female patients (**b**; both sertraline and placebo groups included). Baseline circRNAx levels in SERT-R and SERT-NR patients belonging to a racial minority (c; African American, Asian, and Pacific Islanders). For b-d *p < 0.05, based on two-tailed Mann-Whitney test. Each graph is shown as Mean + SEM with the number of individual biological samples included within each graph.

**Supplementary Figure 2: CircRNAx is particularly enriched in human brain.**

Graph indicates circRNAx (**a**) and circCRY2 (**b**) levels based on circRNA-specific qPCR in total RNA from multiple human organs (brain, heart, lung, kidney, placenta and liver) and cell lines (HelaS3, lymphoblast K-652 and Raji cells). The very highly expression of circRNAx in the human brain tissue is clearly demonstrated. For (a) and (b): *p < 0.05, **p < 0.01, ***p < 0.001, ****p < 0.001, based on an ordinary one-way ANOVA with post-hoc Dunnett's multiple comparisons test (vs Brain). Each graph is shown as Mean + SEM with individual biological samples values included within each graph.

**Supplementary Figure 3: CircRNAx levels in human leukocytes can specifically predict remission following treatment with SSRIs**.

Schematic of the experimental design (**a**). Leukocytes were isolated from human whole blood as described in Materials and Methods and circRNAx expression was quantified with qPCR. Baseline leukocyte circRNAx expression levels in patients that either achieved remission (SSRI-R) or not (SSRI-NR) following 30 weeks of SSRI treatment (**b**). ROC analysis curve for baseline circRNAx levels in leukocytes between SSRI remitters and non-remitters (**c**). The AUC (Area Under the Curve) along with the table of statistics is included. (D Baseline leukocyte circRNAx expression levels in patients that either achieved remission (R) or not (NR) with other classes of antidepressants (Ads) following 30 weeks of treatment (**d**). ADs included SNRIs, TCAs and MAOIs. Baseline circRNAx levels in leukocytes from patients with MDD compared to Bipolar Disorder and unaffected healthy Controls (**e**). For (b): *p < 0.05, based on one sample t-test. Ordinary one-way ANOVA with Tukey's multiple comparisons test was used in (d).

**Supplementary Figure 4: Expression levels of circTulp4 across various mouse brain regions after different *in vivo* pharmacological experiments and circRNAx levels in putamen following 5-HT2A and D2 receptor inhibition.**

Mouse brain circRNAx (**a**) and circTulp4 (**b**) levels after treatment with a pure 5T2AR antagonist MDL100907 and the pure D2R antagonist Sulpiride in mouse putamen. Mouse brain circTulp4 levels after treatment with a pure 5T2AR antagonist MDL100907 and the pure D2R antagonist Sulpiride in mouse frontal cortex (**c**) and nucleus accumbens (**d**). Mouse brain circTulp4 levels after treatment with MK801, a selective NMDAR antagonist and CDPPB, an mGluR5 potent agonist in mouse frontal cortex (**e**) and nucleus accumbens (**f**). **p < 0.01, ***p < 0.001, based on an ordinary one-way ANOVA with post-hoc Dunnett's multiple comparisons test (c, e) or Kruskal Wallis ANOVA with Dunn's multiple comparisons test (a, b). Each graph is shown as Mean + SEM with individual biological samples values included within each graph.

**Supplementary Figure 5: circRNAx and circTulp4 mouse levels in cortical neurons following *in vitro* PKA and PKC inhibitor treatments.**

Primary mouse cortical neuron circRNAx (**a**) and circTulp4 (**b**) levels, based on qPCR, and after treatment with PKA and PKC (small molecule kinase inhibitors). One-way ANOVA with post-hoc Dunnett's multiple comparisons test was used for both (a) and (b). Each graph is shown as Mean + SEM with individual biological samples values included within each graph.

**Supplementary Figure 6: Proposed schematic of circRNAx regulation within neurons**

5-HT2A receptors are found on the postsynaptic membrane of neurons and are G-protein coupled receptors that can activate (among others) the PLC- IP3 pathway, thus leading to the release of Ca+2 from the Endoplasmatic Reticulum (ER). This elevation of Ca+2 levels can activate the downstream protein kinase MEK, which in turn will phosphorylate ERK1/2, resulting in activation of CREB. Tropomyosin receptor kinase B (TrKB) is a transmembrane receptor protein that binds to Brain-derived neurotrophic factor (BDNF). Upon activation of this receptor, autophosphorylation can occur with activation of a series of downstream intracellular kinases (such as Ras-Raf, Akt, PI3K), which can also result in elevation of intracellular Ca+^2^ levels. In a common pathway to 5-HT2A receptor activation, such increases in Ca+^2^ levels can phosphorylate MEK and ERK1/2, thus leading to further CREB activation. CREB-mediated transcriptional effects of yet unknown gene targets then result in the synthesis of circRNAx within the brain.
